# Viral metagenomics of synanthropic urban bats: a surveillance strategy for uncovering potentially zoonotic viruses

**DOI:** 10.1101/2025.07.30.666427

**Authors:** Juliana Amorim Conselheiro, Filipe Romero Rebello Moreira, Gisely Toledo Barone, Adriana Araujo Reis-Menezes, Adriana Rückert da Rosa, Débora Cardoso de Oliveira, Bárbara Aparecida Chaves, Vanderson de Souza Sampaio, Felipe Rocha, Marco Antonio Natal Vigilato, Rodrigo Guerino Stabeli, Rodrigo Fabiano do Carmo Said, Gabriel Luz Wallau, Paulo Eduardo Brandão, Anderson Fernandes de Brito

## Abstract

Bats are natural reservoirs for diverse viruses, including coronaviruses, filoviruses, and paramyxoviruses, many leading to zoonotic spillover events and demanding an integrated surveillance. Here, we present a framework that leverages Brazil’s rabies passive surveillance program to detect bat-borne viruses. Using an algorithm to select representative specimens from 2,422 bats collected across São Paulo state, we submitted 150 paired lung and intestine samples to nanopore metagenomic sequencing. We detected 98 viral contigs from 12 families of public health relevance, including *Arenaviridae, Coronaviridae*, and *Paramyxoviridae*. Notably, the approach identified a previously unknown filovirus in bats in the Americas, validating the framework’s capacity for epidemic preparedness. These findings reveal an undetected viral diversity and demonstrate how existing animal surveillance can monitor pathogen threats. Crucially, in a workshop involving multisectoral One Health experts in Brazil, this framework was validated as a scalable model for national expansion, adapted for low- and middle-income countries (LMICs).

**One-sentence summary line:** The study introduces a viral surveillance method using metagenomics in bats, offering a scalable, cost-effective model for operationalizing One Health surveillance, especially relevant to low- and middle-income countries.

## Introduction

Approximately 75% of emerging infectious diseases over recent decades have zoonotic origins, with the majority involving wildlife reservoirs (Mackenzie et al. 2016). Among these, bats stand out as significant hosts due to their exceptional viral diversity (Letko et al. 2020). To date, over 32,000 viral sequences linked to bats have been deposited in NCBI GenBank, around 88% of which belong to RNA viruses. These include members of families such as *Coronaviridae* (e.g., SARS, MERS), *Rhabdoviridae* (e.g., *Lyssavirus rabies*s), and *Paramyxoviridae* (e.g., Hendra and Nipah viruses) (Van Brussel and Holmes 2022).

Bats represent nearly 25% of all known mammalian species and inhabit a wide range of ecosystems globally (Festa et al. 2023). In Brazil, the order Chiroptera includes nine families, 68 genera, and 186 species (Garbino et al. 2024), of which at least 84 have adapted to urban environments (Nunes et al. 2017). In cities such as São Paulo, members of the Phyllostomidae, Molossidae, and Vespertilionidae families are commonly encountered (Almeida et al. 2015). Synanthropic species, those that thrive in close association with human environments, are particularly relevant to public health surveillance. These animals harbor diverse viruses asymptomatically, and their presence in urban settings increases the likelihood of zoonotic transmission. In this context, the human–bat interface represents a dynamic frontier for emerging infectious diseases.

Despite the recognized risks, viral surveillance in bat populations remains limited. Traditional methods such as polymerase chain reaction (PCR), loop-mediated isothermal amplification (LAMP), and enzyme-linked immunosorbent assays (ELISA) are widely used for pathogen detection but are typically restricted to known viruses (Kawasaki et al. 2023). In contrast, metagenomic sequencing offers a powerful alternative for unbiased viral discovery, capable of identifying divergent or novel viruses in a single assay (Miller et al. 2013; Rahimian and Panahi 2024). Although cost and computational demands remain as challenges, the broad-spectrum nature of metagenomic data presents significant advantages.

In Brazil, the Laboratory of Diagnostics of Zoonosis and Vector-Borne Diseases (LabZoo, DVZ/COVISA/SEABEVS/SMS/PMSP) receives approximately 3,000 bats per year via the city’s rabies passive surveillance program. Passive surveillance, particularly for wildlife, refers to specimen collection initiated by citizen or health authority reports rather than active field sampling (Minter et al. 2024). While such programs may introduce sampling biases, they represent cost-effective platforms for detecting circulating or emerging pathogens in large urban areas (Gilbert and Cliffe 2016).

Despite the availability of these specimens, there is currently no integrated strategy to leverage them for broad viral discovery or early zoonotic risk detection. As global health increasingly embraces the One Health paradigm, utilizing urban wildlife surveillance for early viral detection is both a practical and necessary innovation. To address this gap, we developed a novel, cost-effective metagenomic surveillance framework tailored to passive rabies surveillance samples. Our strategy combines an objective sample selection algorithm with SMART-9N/Nanopore sequencing and a robust bioinformatics pipeline, applied to paired lung and intestinal tissues from synanthropic bats collected in São Paulo State, Brazil.

## Results

A total of 312 (13%) of all 2,422 bats received at LabZoo for rabies diagnostics through passive surveillance were found eligible for viral metagenomic sequencing, from which 150 specimens with the highest scores (>4 points) were selected for sequencing (6.19% - 150/2,422), totaling 300 samples (intestines and lungs separately). The samples came from 14 municipalities within the State of São Paulo, Brazil (**Fig. 1A; B**), which together account for over 17.3 million inhabitants and have an average population density of approximately 3,220 inhabitants/km^2^.

**Figure 1.**
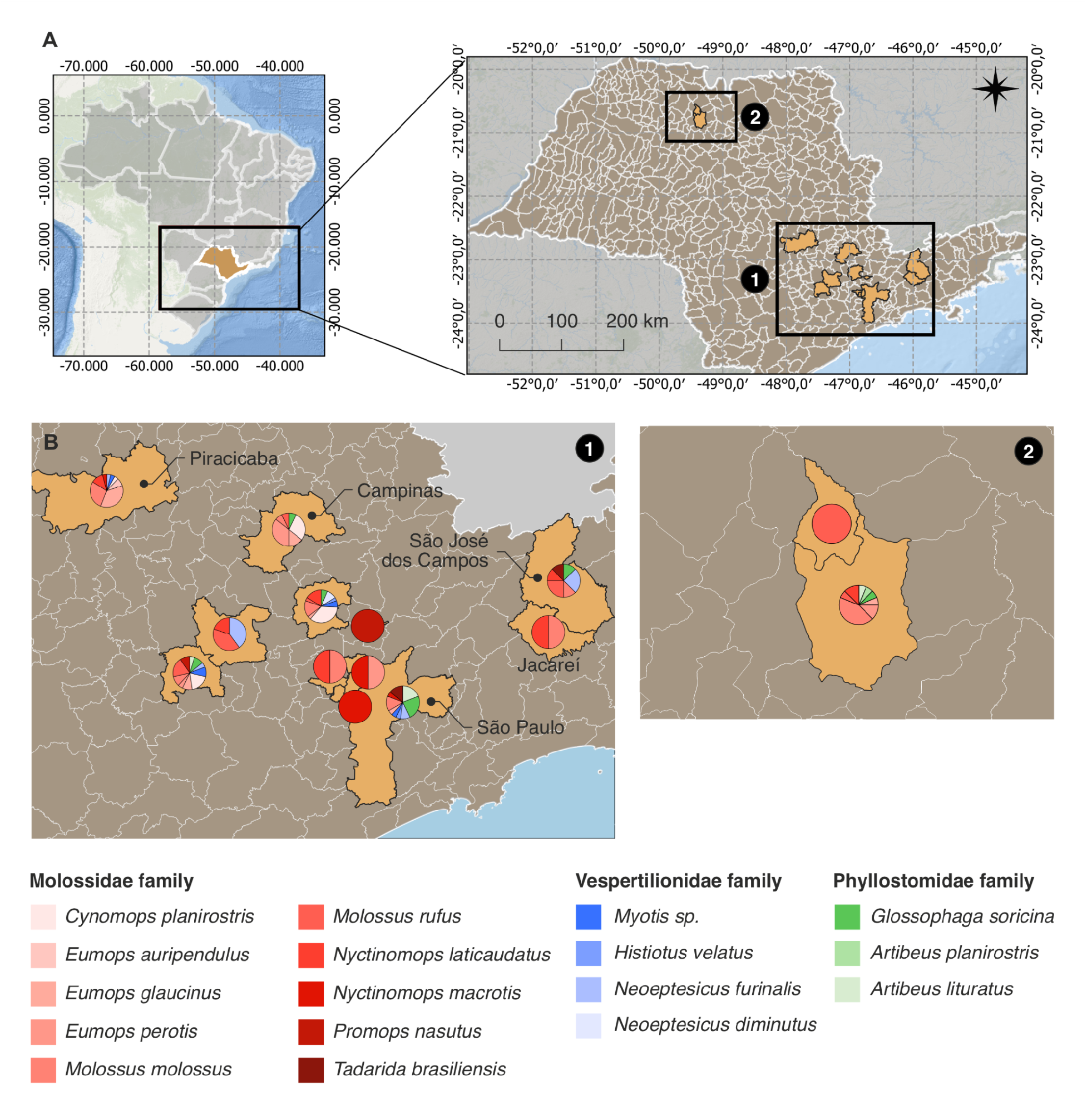
Origin of the 150 bats submitted to viral metagenomics. **A**. A total of 14 municipalities within the State of São Paulo, Brazil, are represented. **B**. Colored circles within each municipality represent the bat species.

The bat species belonged to three families: Molossidae (n = 112 specimens), Phyllostomidae (n = 19 specimens), and Vespertillionidae (n = 19 specimens). The greatest diversity was found amongst the Molossidae family, being *Molossus molossus* the most represented species (n = 28) (**Fig. 2A**). As recommended by the rabies surveillance system, bats found near human dwellings (dead or alive) are constantly reported by local citizens. As such, the majority of the specimens included in this study were found in that context. The circumstances under which selected bats were collected were classified according to the standardized categories used by DVZ (detailed definitions are available at Methods section) (Fig. 2B). A total of 40% (60/150) of the specimens were found on the ground, while 10 (15/150) and 9.3% (14/150) were found inside houses or in external areas of residences, respectively. Smaller proportions were found in roosts (5/150, 3.3%), hanging (3/150, 2%) or were brought in by pets (2/150, 1.3%). For 34% (51/150), no information on the circumstances of collection was reported. Contact with humans or pets was reported for 25% (37/150).

**Figure 2.**
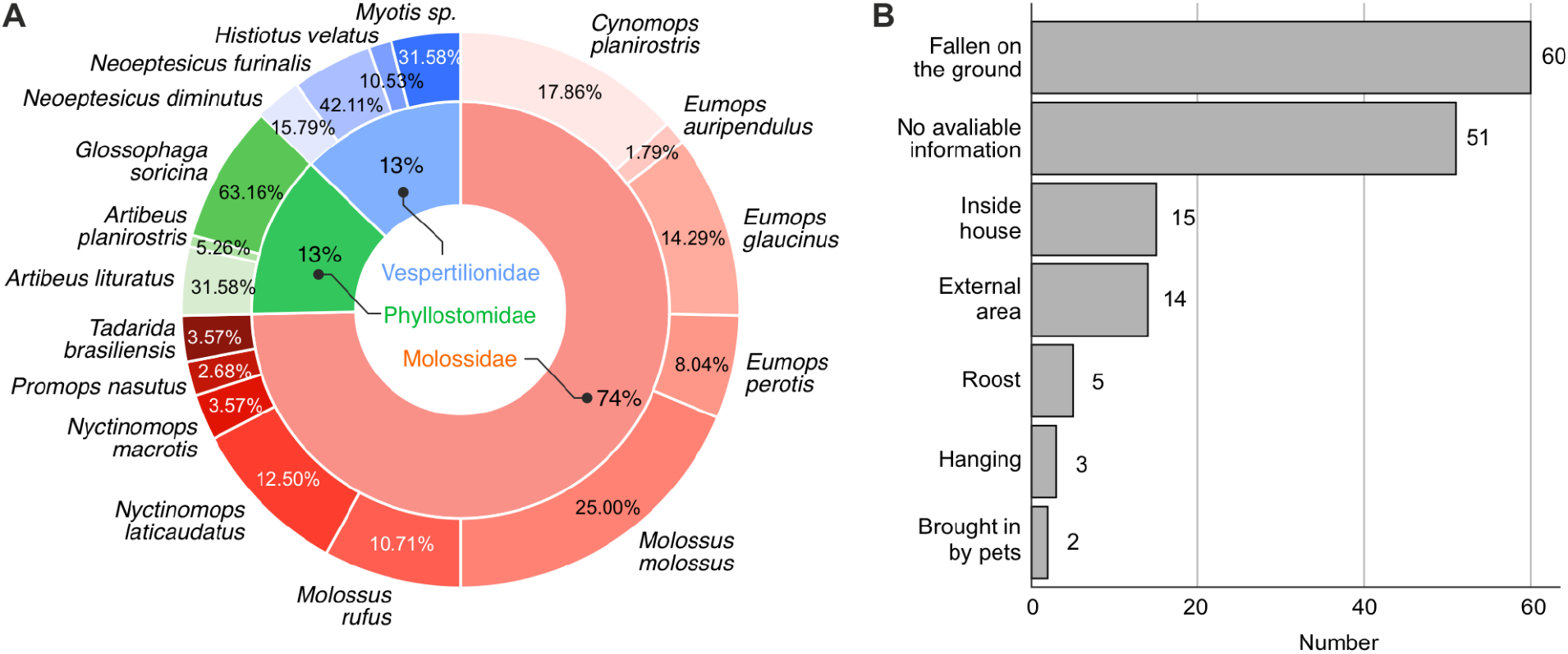
Distribution of bat species and circumstances of collection. **A**. Distribution of bats species used in viral metagenomics, categorized by family: Molossidae, Phyllostomidae, and Vespertillionidae. **B**. Circumstances under which bat carcasses were found as part of passive surveillance submissions. Definitions of each category are provided in the Methods section.

### Metagenomics analysis of bats collected through passive rabies surveillance enables detection of viruses with zoonotic potential

A total of 34.3 million nanopore reads were generated in this study, from which 88.2% passed quality control **(Table 1**). A total of 3,260 (approximately 0.01%) of these reads were mapped against *de novo* assembled viral contigs (n = 171), spanning 18 distinct viral families. A complete list of these contigs is available on **Supplementary Table 1**. To ensure the reliability of the viral assignments, we applied a series of minimum filters to avoid artifactual detection of viruses, including the establishment of coverage thresholds, index-hopping filters and secondary similarity searches against a broader database (see Methods). After these filtering steps, 114 contigs remained (**Tables 1** and **2**), 66 assembled from intestines samples and 48 from lung samples, representing diverse viral families, including *Arenaviridae, Chuviridae, Coronaviridae, Endornaviridae, Filoviridae, Flaviviridae, Herelleviridae, Paramyxoviridae, Peribunyaviridae, Permutotetraviridae, Phenuiviridae, Picornaviridae, Poxviridae, Retroviridae, Rhabdoviridae, Sedoreoviridae, Solemoviridae* and *Virgaviridae*. A total of 16% (24/150) bats showed viral hits in at least one of the viscera analyzed, comprehending 10 distinct taxa, such as *Artibeus lituratus, Neoeptesicus furinalis, Eumops glaucinus, Eumops perotis, Glossophaga soricina, Histiotus velatus, Molossus molossus, Molossus rufus, Myotis sp*. and *Nyctinomops laticaudatus* (**Fig. 3**).

**Table 1.** Metrics obtained by viral metagenomics of 150 bat from São Paulo State, Brazil.

| Viscera | Total raw reads | Total trimmed reads | Total viral reads | Total viral contigs | Filtered viral reads* | Filtered viral contigs* |
| --- | --- | --- | --- | --- | --- | --- |
| Lungs | 22,860,735 | 20,613,332 | 768 | 70 | 743 | 48 |
| Intestines | 11,471,903 | 9,679,906 | 2,492 | 101 | 2,439 | 66 |
| **TOTAL** | **34,332,638** | **30,293,238** | **3,260** | **171** | **3,182** | **114** |

\*Sequences were filtered as described in the Methods section.

**Figure 3.**
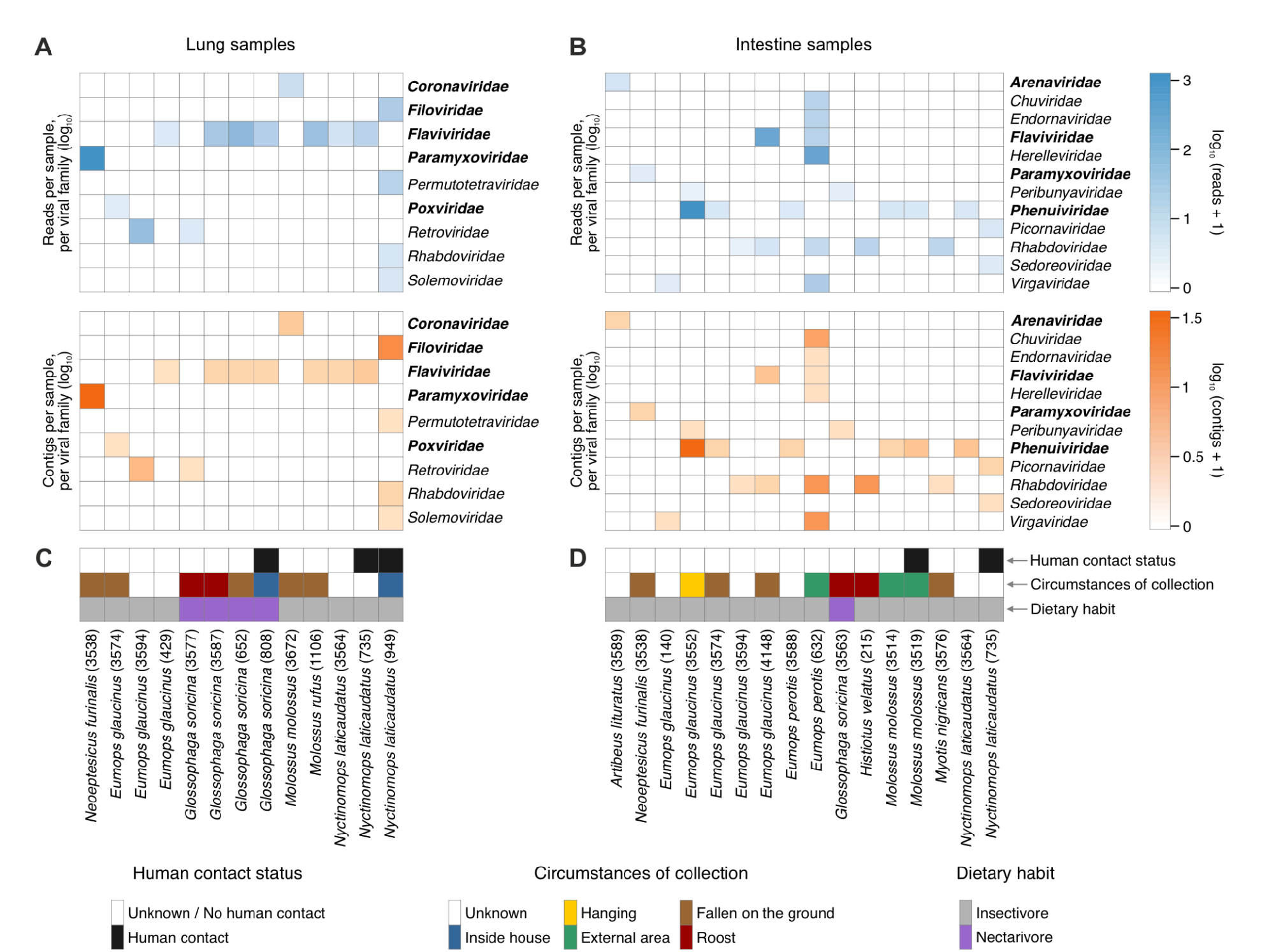
Tissue-specific viral signals in bat metagenomes. Heatmaps show (top) the number of viral contigs and (bottom) read abundance (total reads mapped to contigs assigned to each family) for each bat sample. Results are displayed separately for lungs (**A**) and intestines (**B**). Values are plotted on a log_10_ scale. Viral families that include pathogens prioritized by the World Health Organization R&D Blueprint as having high potential for causing public-health emergencies of international concern are indicated in bold. Panels (**C**) and (**D**) show information about the history of human contact, circumstances of collection and dietary habits of the sequenced bat specimens.

In the lungs, we identified 48 viral contigs primarily classified into nine families: *Coronaviridae* (genus *Alphacoronavirus*, n = 3), *Filoviridae* (genera *Orthoebolavirus*/*Cuevavirus*, n = 8), *Flaviviridae* (genera *Pegivirus* and *Hepacivirus*, n = 14), *Paramyxoviridae* (genus *Morbillivirus*, n = 13), *Permutotetraviridae* (n = 1), *Poxviridae* (genus *Avipoxvirus*, n = 1), *Retroviridae* (genus *Betaretrovirus*, n = 5), *Rhabdoviridae* (genus *Lyssavirus*, n = 2), and *Solemoviridae* (n = 1) (**Fig. 3A**; **Table 2**; **Supplementary Table 1**). *Flaviviridae* was the most frequently detected family, occurring in seven samples from *E. glaucinus, G. soricina, M. rufus* and *N. laticaudatus*, followed by *Retroviridae*, detected in two samples from *E. glaucinus* and *G. soricina*. All other families were found in a single sample from different host species, including *E. furinalis* (*Paramyxoviridae*), *E. glaucinus* (*Poxviridae*), *M. molossus* (*Coronaviridae*) and *N. laticaudatus* (*Filoviridae, Permutotetraviridae, Rhabdoviridae* and *Solemoviridae*). In addition to these viruses with potential zoonotic relevance, from priority viral families, according to the World Health Organization R&D Blueprint (WHO 2024), we also highlight the detection of a filovirus in bats in the Americas.

In the intestines, we identified 66 viral contigs spanning 12 families across 24 samples (Fig. 3A). Among viral families of public-health relevance, we detected *Flaviviridae* (n = 4), *Paramyxoviridae* (n = 2), *Peribunyaviridae* (n = 2), *Phenuiviridae* (n = 23), *Picornaviridae* (n = 2), *Rhabdoviridae* (n = 16), and *Sedoreoviridae* (n = 1). *Phenuiviridae* was the most common family, represented by 23 contigs from six samples. We also detected families typically associated with non-mammalian hosts — *Chuviridae* (n = 5), *Endornaviridae* (n = 1), *Herelleviridae* (n = 1), and *Virgaviridae* (n = 7) (**Fig. 3B**; **Table 2**; **Supplementary Table 1**). While some taxa likely represent true infections — such as *Mammarenavirus* (*Arenaviridae*) in *A. lituratus, Morbillivirus* (*Paramyxoviridae*) in *E. furinalis, Shanbavirus* (*Picornaviridae*) and *Rotavirus* (*Sedoreoviridae*) in *N. laticaudatus* — several others are more plausibly of dietary or environmental origin (*e*.*g*., derived from insects, plants, or phages). These include phenuiviruses (genera *Phlebovirus, Ixovirus, Uukuvirus*), some rhabdoviruses (genera *Alpha*- and *Betanemrhavirus, Gammaplatrhavirus*), and members of *Chuviridae, Endornaviridae, Herelleviridae, Peribunyaviridae* and *Virgaviridae*.

Although three viral families were detected in both lungs and intestines across all samples (*Flaviviridae, Paramyxoviridae*, and *Rhabdoviridae)* only *Morbillivirus* (*Paramyxoviridae*) was detected in both tissues from the same animal. Genus-level assessment indicates that different viral groups were associated with each tissue for the other families: *Hepacivirus* and *Pegivirus* occurred in lungs, whereas *Orthoflavivirus* members were found in intestines; similarly, *Lyssavirus* was detected in lungs, while other rhabdovirus genera occurred only in intestines. Because several of these families include viruses of public-health concern, we also evaluated the circumstances under which infected bats were collected. Infected individuals were obtained across a range of situations, and 4/16 (25%) had reported contact with humans, including bats carrying members of *Filoviridae, Flaviviridae, Picornaviridae, Rhabdoviridae*, and *Sedoreoviridae* (**Fig. 1**; **Table 2**).

### Phylogenetic analyses clarify the evolutionary placement of novel viral sequences

To contextualize the novel sequences identified from viral families of particular zoonotic relevance, we performed maximum-likelihood (ML) phylogenetic reconstructions using comprehensive reference datasets. Overall, these analyses were consistent with the known evolutionary history of each viral family and aligned with preliminary classifications based on sequence similarity searches. These reconstructions provided refined insights into the evolutionary placement of the novel sequences relative to the closest available references (**Fig. 4**). All multiple sequence alignments and phylogenetic trees generated in this study are provided in **Supplementary File 1**.

**Figure 4.**
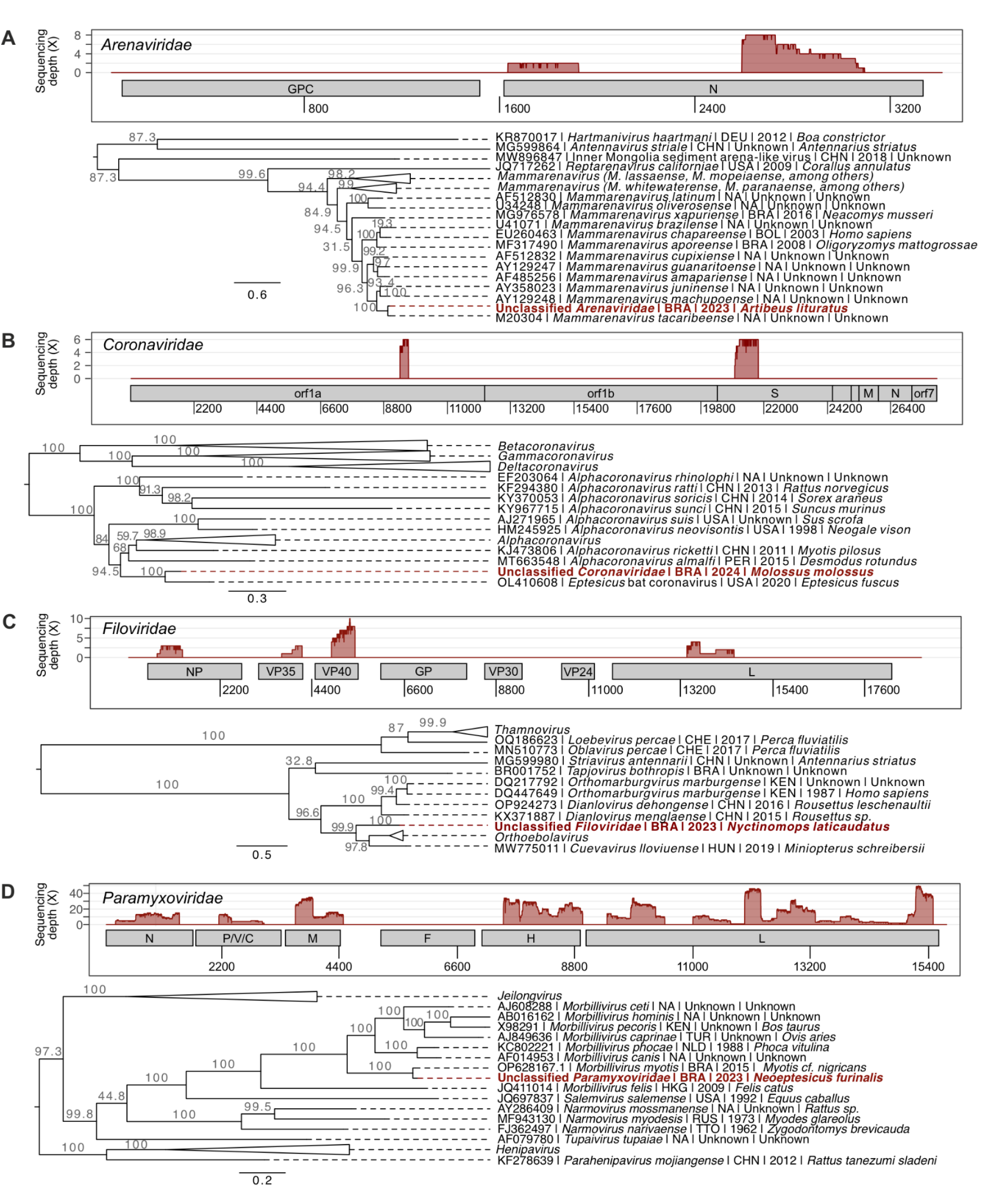
Read mapping and maximum-likelihood phylogenetic analyses of amino acid sequences translated from coding regions within assembled scaffolds assigned to viral families of zoonotic relevance. For each panel, sequencing depth and read mapping to a reference sequence are shown, followed by phylogenetic inference of the corresponding protein regions, with tip labels in red indicating sequences generated in this study. **(A)** Reads mapped to a *Mammarenavirus tacaribense* segment S reference sequence (ON648821), followed by a phylogeny of nucleoprotein sequences from members of *Arenaviridae*. **(B)** Reads mapped to an *Eptesicus* bat coronavirus reference genome (OL410608), followed by a phylogeny based on concatenated *orf1a* and *S* protein fragments from members of *Coronaviridae*. **(C)** Reads mapped to an *Orthoebolavirus zairense* reference genome (AF086833), followed by a phylogeny based on concatenated NP, VP35, VP40, and L protein fragments from members of *Filoviridae*. **(D)** Reads mapped to a *Morbillivirus myotis* reference genome (OP628167), followed by a phylogeny based on concatenated fragments spanning most encoded proteins from members of *Paramyxoviridae*. Tip labels show GenBank accession numbers, viral species, country of origin (ISO 3166 alpha-3 code), sample collection year, and the host species from which the sequences were derived. Scale bars represent amino acid substitutions per site, and collapsed clades indicate sequences belonging to the same genus-level taxa. Complete phylogenetic trees are provided in **Supplementary File 1**.

The unclassified *Arenaviridae* scaffold sequence (791 bp), detected in an intestine sample from an *Artibeus lituratus* specimen, showed highest similarity to a *Mammarenavirus tacaribense* based on tBLASTn results (95.7% identity, 91% query coverage, e-value: 1e-103). This classification was corroborated by ML phylogenetic analysis (**Fig. 4A**), where the sequence clustered within a clade of a previously characterized *M. tacaribense* isolate (SH-aLRT = 100), and other *Mammarenavirus* isolates collected between 2003 and 2016 in Brazil, Bolivia, and other countries.

The unclassified *Coronaviridae* scaffold sequence (1128 bp), detected in a lung sample from a *Molossus molossus* specimen, showed highest similarity to an unclassified *Alphacoronavirus* found in a *Eptesicus fuscus* bat (tBLASTn results: 80.9% identity, 85% query coverage, e-value: 9e-50). Phylogenetic analysis supported this genus-level classification: the novel sequence formed a sister clade (SH-aLRT = 94.5) to a group of sequences derived from other viruses falling within the *Alphacoronavirus* genus (**Fig. 4B**). These reference sequences originated mostly from Vespertilionidae bats sampled between 2005 and 2020 in China, Peru, the United States and other countries.

The unclassified *Filoviridae* scaffold sequence (2802 bp), detected in a lung sample from a *Nyctinomops laticaudatus* specimen, had as its closest match sequences from *Orthoebolavirus restonense* (tBLASTn results: 72.8% identity, 100% query coverage, e-value: 3e-178). Phylogenetic analysis indicated that the novel sequence clusters externally to filoviruses belonging to the genera *Orthoebolavirus* and *Cuevavirus*, sampled from humans and bats collected between 1994 and 2019 in Africa and Europe (**Fig. 4C**). As the formal classification of novel filoviruses requires complete genome sequences, the precise taxonomic status of this virus remains unresolved. Nonetheless, to our knowledge, this represents the first detection of a filovirus in bats from the Americas.

The unclassified *Paramyxoviridae* scaffold sequence, detected in a lung sample from a *Neoeptesicus furinalis* specimen, was most similar to that of *Morbillivirus myotis* (tBLASTn results: 89.1% identity, 91% query coverage, e-value: 0). This identification was further supported by phylogenetic analysis, where the sequence clustered within a clade of *Morbillivirus* sequences (SH-aLRT = 100), grouping with a sequence previously sampled in Brazil in 2015 (**Fig. 4D**).

## Discussion

Emerging infectious diseases with zoonotic origins have repeatedly demonstrated the need for scalable, pre-emptive viral surveillance in wildlife, especially in synanthropic species that frequently interface with human environments. In this study, we present a metagenomics-based surveillance framework applied to bats sampled through Brazil’s passive rabies monitoring system. This approach leverages existing infrastructure to cost-effectively detect viruses with zoonotic potential, including families of high public health concern such as *Filoviridae, Paramyxoviridae*, and *Coronaviridae*. Notably, we report what is, to our knowledge, the first detection of a filovirus in bats in the Americas. These findings have translated into immediate policy dialogue: the framework developed here was evaluated in February 2025, in a technical workshop hosted by the Pan American Health Organization (PAHO/WHO), involving the Brazilian Ministry of Health, National Councils of State and Municipal Health Secretariats (CONASS and CONASEMS) and multisectoral experts from the animal and environmental health sectors. In that occasion, this framework was recognized as a technically robust and operationally viable model to guide the implementation of a national surveillance protocol (PAHO 2026, 2025).

As bats are legally protected in many countries, including Brazil, and are difficult to sample longitudinally in the wild, this community-based passive surveillance framework offers an ethical and logistically feasible alternative for accessing specimens (Minter et al. 2024). Although its effectiveness can vary with disease dynamics and public reporting patterns, our findings support its value for virus detection and discovery. This premise was validated by One Health experts in Brazil, who agreed that leveraging the flow of bat samples from passive rabies surveillance constitutes a viable, practical, and replicable strategy for expanding genomic surveillance to other regions (PAHO 2026, 2025). At LabZoo, more than 3,000 bats are submitted annually for rabies diagnostics, yet these specimens are typically not screened for other pathogens. Our metagenomics pipeline demonstrates that this underutilized resource can be mined for broad viral surveillance.

Due to the high cost of metagenomic processing, we prioritized tissues more directly associated with spillover potential, *i. e*. lung and intestinal tissues. The intestine represents a major interface between the host and the external environment, as viral particles shed through feces can contaminate habitats or be transmitted to other mammals, including humans (Temmam et al. 2014).

Conversely, the lung is the primary site for respiratory infections and is capable of initiating cross-species viral spillover (He et al. 2022; Wu et al. 2021). Therefore, both organs offer valuable insights into potential spillover routes. Other organs, such as liver and spleen, were not included in this analysis due to their lower direct relevance to transmission pathways in the context of surveillance-oriented studies.

We found a distinct separation of viral families between intestinal and pulmonary tissues. The aggregated intestinal virome predominantly harbored *Phenuiviridae*, a diverse group with members capable of infecting arthropods, plants, and vertebrates, including Rift Valley fever virus and Severe Fever with Thrombocytopenia Syndrome virus (Sasaya et al. 2023), whereas the lungs featured more vertebrate-related viruses, including *Arenaviridae* (genus *Mammarenavirus*), *Paramyxoviridae* (genus *Morbillivirus*), *Coronaviridae* (genus *Alphacoronavirus*), and, at a higher frequency, *Flaviviridae* (genus *Pegivirus*). These patterns suggest that tissue tropism, dietary exposure, and possibly arthropod infestation contribute to virome composition (Van Brussel and Holmes 2022; Bazzoni et al. 2024). These findings support incorporating multi-tissue sampling into future viral metagenomics workflows, in agreement with previous recommendations (Wallau et al. 2023).

The presence of tick-associated viruses in intestines, particularly members of *Phenuiviridae* (**Fig. 3B**), may reflect ectoparasite ingestion or infestation rather than active infection, consistent with reports of tick-borne viruses associated with bats (Xu et al. 2022; Luz et al. 2016). This raises intriguing ecological questions about the role of ectoparasites in bat viral ecology that merit further investigation. The detection of *Flaviviridae* sequences across multiple samples (**Fig. 3**) is also consistent with the literature, which indicates that bats are natural hosts frequently infected with members of the genera *Hepacivirus* and *Pegivirus*, typically without apparent disease (Quan et al. 2013).

Among the vertebrate-associated viruses, we recovered a near-complete genome of a *Myotis* bat morbillivirus, previously identified in Brazil and shown to recognize human cell receptors *in vitro* (**Fig. 4D**) (Wells et al. 2022; Ikegame et al. 2023). Given the known capacity of paramyxoviruses to cross species barriers (Branda et al. 2024), this detection underscores the need for continuous surveillance. We also detected a sequence closely related to *Mammarenavirus tacaribeense* (**Fig. 4A**), which has been reported in bats and ticks (Bentim Góes et al. 2022; Sayler et al. 2014), suggesting possible multi-host dynamics. Additionally, the detection of a new alphacoronavirus in *Molossus molossus* aligns with recent findings of new coronaviruses in the Americas (Cerri et al. 2023; Schaeffer et al. 2022; Wallau et al. 2025), shedding more light into the growing diversity of bat-borne coronaviruses in the region.

The most striking finding of this study was the identification of sequences of an unclassified *Filoviridae*, which represents the first evidence of a filovirus in bats from the Americas (**Fig. 4C**). In the initial sequencing round, a single read matching *Filoviridae* was detected in the lung sample of one individual (specimen 949, *Nyctinomops laticaudatus*). To verify this unexpected result, the same specimen was reprocessed, and a second metagenomic sequencing run was performed using freshly extracted material from the lung, intestine, and also the liver. The resequencing confirmed additional *Filoviridae*-related reads exclusively in the lung, while both intestinal and hepatic samples resulted negative. This consistency indicates that the viral signal was reproducible and restricted to respiratory tissue, reinforcing the relevance of lung sampling for detecting potential zoonotic viruses. Nevertheless, given the limited number of reads, this observation requires further validation through targeted PCR, deeper sequencing, or screening of additional specimens.

Despite its important findings, our study has some limitations. The most important relates to the variable time interval between the death of the animals and the collection of internal organs, which was beyond the control of the proposed framework, as specimens were obtained through passive surveillance. This likely affected RNA preservation, since post-mortem degradation can occur rapidly under field conditions, potentially reducing the sensitivity of viral detection, contributing to the low sequencing depth and coverage, and resulting in incomplete genome assemblies for many detected viruses. Additional limitations include the lack of PCR or culture-based confirmation. Going forward, integrating targeted PCR assays, virus isolation, and host-receptor interaction studies will be critical to characterize the biological properties of these viruses. We also recommend automation of the decision-tree selection algorithm and integration of ecological metadata to optimize the detection of viruses with zoonotic potential. Finally, to ensure the sustainability and impact of this surveillance model, recommendations have been made to include bat viral surveillance in the agenda of the National One Health Committee, fostering intersectoral governance and the development of a standardized national protocol (PAHO 2026, 2025).

## Conclusion

Our findings demonstrate that viral metagenomics applied to bat specimens from a rabies passive surveillance system can reveal viruses of zoonotic relevance, including novel or divergent lineages. Following a technical evaluation by Public Health authorities in Brazil, this approach was validated as a scalable and cost-effective model for operationalizing One Health surveillance, with potential applications in pandemic prevention, particularly in low- and middle-income countries.

Furthermore, the operational success of this framework has catalyzed policy action, resulting in a formal recommendation to include bat viral surveillance in the permanent agenda of the National One Health Committee to ensure institutional sustainability (PAHO 2026, 2025). Findings like the ones reported in this study are relevant for triggering active surveillance of novel viruses, especially when suspected or confirmed human cases of new zoonotic diseases are reported. As ecological disruption continues to drive cross-species transmission events, strategies like the one proposed here will be critical for the early detection of zoonotic viruses with epidemic and pandemic potential.

## Methods

### Sample collection and selection protocol

Bats were obtained through passive surveillance activities for rabies monitoring and were received at LabZoo already deceased. Information about the circumstances in which carcasses were found was recorded whenever available. Each specimen was classified according to standardized categories used by DVZ as follows: “Fallen on the ground” refers to bats found lying on the ground in public or private areas, often reported by citizens; “Inside houses” denotes carcasses or moribund animals recovered inside human dwellings, while “External Area” refers to animals collected in yards, gardens, or other peridomestic outdoor environments; “Roost” indicates animals obtained directly from synanthropic roosting sites such as roofs or abandoned buildings; “Hanging” refers to bats found suspended from walls, eaves, or vegetation but already dead or near death; “Brought in by pets” includes animals that were preyed upon by domestic animals and carried into houses; Finally, “No available information” was applied when the submitter did not provide any contextual details. This classification was used to contextualize the origin of samples and evaluate potential environmental or epidemiological factors associated with passive surveillance submissions.

Upon arrival, carcasses were stored at –20 °C until taxonomic identification and brain tissue was collected for rabies diagnosis following the gold standard procedure which is direct immunofluorescence assay. Following these procedures, viscera (lung and intestine) were aseptically removed and immediately stored at –80 °C until viral metagenomic processing. These organs were selected because they represent key interfaces for potential zoonotic transmission, such as respiratory and enteric routes, while also providing high viral diversity (He et al. 2022; Wu et al. 2021)(Temmam et al. 2014). All tissue manipulations were conducted under appropriate biosafety conditions within a Class II biological safety cabinet. Although the original purpose of sample collection was rabies diagnosis, the use of these specimens in the present study complies with the Brazilian Ministry of Health’s Ordinance No. 1138. Article 3 of this ordinance authorizes Zoonoses Surveillance Units to: receive animal carcasses (item XI), perform laboratory diagnostics for zoonoses of public health relevance (item IV), and collect biological samples for diagnostic purposes beyond rabies (item XIV). The samples originated from 22 municipalities of São Paulo State, Brazil, between October 2023 and February 2024. A decision tree was elaborated to guide selection of samples (**Fig. 5**; see **Supplementary Methods 1** and **Supplementary Table 2**), with specimens scoring between 5 and 7 deemed eligible for viral metagenomic sequencing.

**Figure 5.**
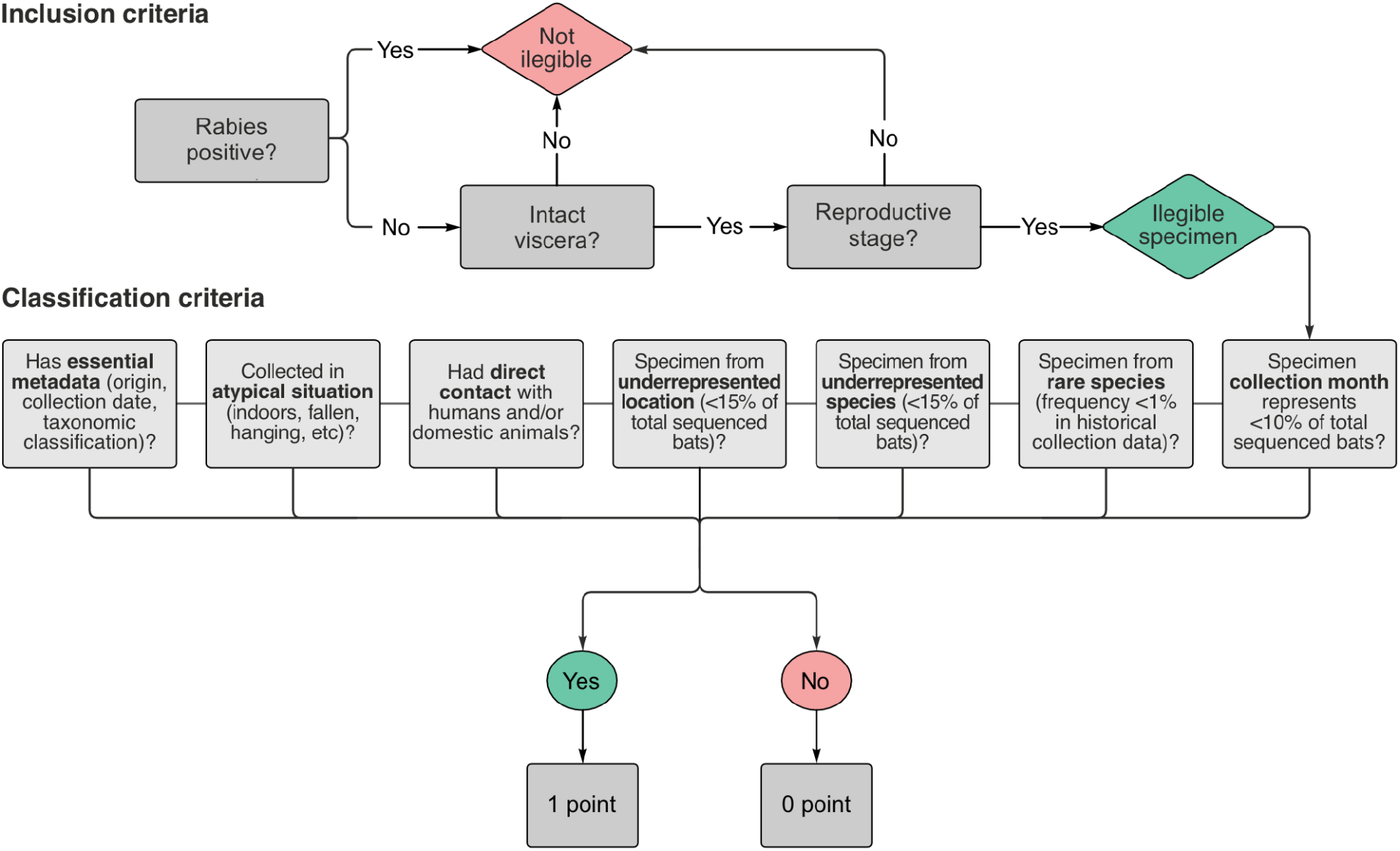
Decision tree used to guide the selection of bats for viral metagenomics. Viscera with vivid red color and visually integral texture were considered intact. Only specimens at reproductive stages (i.e. not infants) were eligible, and those positive for rabies were excluded. Thresholds for municipality representation (<15%), species representation (<5%), rare species (<1%), and collection month (<10%) were calculated using the empirical distribution of previously sequenced samples. Each parameter scores one point whenever its threshold is met. This scoring system was designed to minimize bias and ensure a balanced representation of bats across space, time and species. See **Supplementary Table 2**.

### Bats taxonomic classification

Taxonomic identification of bat specimens was based on external and cranial morphological characteristics using dichotomous keys as described elsewhere (refs). The classification followed the taxonomic framework adopted by DVZ. Identification was carried out by trained personnel and species were assigned according to current standards for Chiroptera taxonomy (Diaz et al. 2021; Reis et al. 2006; Vizotto and Taddei 1973; Cláudio et al. 2023).

### Nucleic acid isolation

A total of 100uL of Phosphate-Buffered Saline pH 7.4 (PBS) along with 4 glass beads of 1mm diameter were added to 10mg of intestine or lung tissue of each bat and then submitted to tissue homogenization in an L-Beader 24 (Loccus, Cotia, SP, Brazil). Next, 500μL of PBS were added to each sample and the tubes were centrifuged at 16,000 x *g* for 5 minutes at 4°C. The supernatants were filtered through 0.45μM syringe filters (Sigma-Aldrich, St. Louis, MO, USA) and submitted to nucleic acid isolation using the Bio Gene Viral DNA/RNA extraction kit (Quibasa, Belo Horizonte, MG, Brazil), following the manufacturer’s instructions. The samples were also submitted to DNAse treatment using TURBO DNAse 2U/μL (Invitrogen, Carlsbad, CA, USA), followed by DNAse cleanup using the RNA Clean & Concentrator-5 kit (Zymo Research, Tustin, CA, USA) as described by each manufacturer’s protocol.

### Portable nanopore sequencing

To enable the implementation of unbiased viral metagenomics within a real-world surveillance setting, we opted for portable nanopore sequencing as the core sequencing strategy. Unlike conventional high-throughput platforms, which require specialized laboratory infrastructure, controlled environments, and substantial operational budgets, nanopore devices offer a compact, low-cost, and field-adaptable alternative that can be deployed within existing public health workflows. This approach was particularly suited to our context, where surveillance activities are conducted in facilities not optimized for advanced molecular genomics. By leveraging portable sequencing, we sought to evaluate the feasibility of generating meaningful viral discovery data under operational constraints and to develop a workflow that could be readily generalized to similar resource-limited surveillance settings. Our aim was not only to characterize bat-associated viral diversity but also to demonstrate that metagenomic approaches can be effectively integrated into routine surveillance systems, even in the absence of ideal laboratory infrastructure. Although this strategy does not yield the depth of coverage necessary for comprehensive virome reconstruction or complete genome assembly, our primary objective was detection, establishing whether portable sequencing could reliably identify viral taxa of public health relevance within the constraints of routine surveillance.

The synthesis of cDNA and PCR reaction were performed using the SMART-9N protocol, as described elsewhere (Claro et al. 2021). PCR products were purified using a 1:1 ratio of AMPure XP beads (Beckman Coulter, High Wycombe, UK), quantified with the fluorometer Qubit 3.0 (Life Technologies, Carlsbad, CA, USA), and normalized to 50ng per sample. Library preparation was carried out using the Ultra II End Repair/dA-tailing Module (NEB) and Ultra Ligation Module (NEB). Barcoding was performed using the EXP-NBD104 and EXP-NBD114 kits along with the Ligation Sequencing kit SQK-LSK109 (ONT, Oxford, UK). Next generation sequencing was carried out in the MinION MK1C (ONT), by applying 50ng of the final libraries onto 14 R9.4.1 flow cells (ONT). A total of 24 samples were sequenced per individual flow cell. Sequencing was performed for 72 hours or until complete pore depletion.

### Bioinformatics workflow and sequence analysis

Raw sequencing reads were basecalled and demultiplexed using Guppy (Oxford Nanopore Technologies, ONT). Quality control analyses were conducted with FastQC v0.12.1 (http://www.bioinformatics.babraham.ac.uk/projects/fastqc/) and pycoQC v2.5.2. Primer and barcode trimming was performed using Cutadapt v4.8 and Porechop v0.3.2 (ONT). We performed sensitivity analysis with three different *de novo* assemblers - Flye, Raven, and MEGAHIT - and determined the latter to be the most sensitive for assembling viral sequences in our dataset (details on **Supplementary Methods 2**).

Assemblies were polished using Medaka v1.11.3 (ONT) to generate high-quality consensus sequences. These sequences were queried against the NCBI RefSeq viral protein database (downloaded on May 10, 2025) using DIAMOND v2.1.9 (Buchfink et al. 2021) in blastx mode with the ultra-sensitive option enabled. To minimize false positives, hits with an e-value greater than 1e-10 were filtered out. For each sample, trimmed reads were remapped to viral contigs using Minimap2 (Li 2018) with parameters optimized for sensitivity (-k10, -w1, -s50). Samtools (Li et al. 2009) and Bedtools (Quinlan and Hall 2010) were used to quantify the number of mapped reads and calculate sequencing coverage, respectively. Sequences primarily assigned to viruses were further analyzed via diamond blastx searches against the NCBI non-redundant (nr) database to exclude potential false positives. Taxonomic criteria were operationalized according to ICTV guidelines for each viral group (family, genus, species). Following previous studies in viral metagenomics (Geoghegan et al. 2021; Asplund et al. 2019), we applied an index-hopping filter at the family level to remove spurious assignments. Specifically, a viral contig was considered valid only if it was supported by a number of reads greater than 0.5% of the maximum read count observed for that family across all samples in the same sequencing run. Also, contigs supported by fewer than two reads were excluded from further analysis. Open reading frames (ORFs) for relevant viral contigs were identified with the NCBI ORFfinder web-app.

### Phylogenetic analyses

Viral genome scaffolds assigned to families of zoonotic relevance were subjected to phylogenetic analyses. Phylogenetic trees were reconstructed using concatenated protein fragments translated from multiple coding regions within the assembled viral scaffolds. Reference datasets were compiled by retrieving protein sequences from reference genomes for each viral family, as compiled by the International Committee on Taxonomy of Viruses - ICTV. Sequence alignments were performed using MUSCLE (Edgar 2004) with default parameters and subsequently trimmed with TrimAl (Capella-Gutiérrez et al. 2009) using the -gappyout option. Maximum Likelihood phylogenetic trees were inferred with IQ-TREE2 v2.0.7 (Minh et al. 2020), under the best-fitting amino acid substitution model determined by ModelFinder (Kalyaanamoorthy et al. 2017). Branch support was assessed using 1,000 replicates of the Shimodaira–Hasegawa approximate likelihood ratio test (SH-aLRT) (Guindon et al. 2010).

### Statistical analysis

To evaluate differences in viral community composition between intestinal and pulmonary samples, we computed pairwise dissimilarities using the Bray-Curtis index, which captures both the presence/absence and relative abundance of viral taxa. Statistical significance of compositional differences between groups was assessed using Permutational Multivariate Analysis of Variance (PERMANOVA), a non-parametric method based on permutations of the distance matrix that tests for differences in multivariate centroid locations among groups (Anderson 2008). A p-value < 0.05 was considered statistically significant. All analyses were performed in R Statistical Software v4.3.1 (R Core Team 2021) using RStudio v2024.04.2. Ecological analyses were conducted using the vegan package v2.6-8 (Oksanen et al. 2001), and data visualization was performed with ggplot2 (Wickham 2016).

## Acknowledgments

This work was supported by Instituto Todos pela Saúde (ITpS), under the project A125. We thank Pan American Health Organization (PAHO) for technical-scientific and logistical support. The PAHO authors alone are responsible for the views expressed in this publication, and they do not necessarily represent the decisions or policies of the Pan American Health Organization.

## Author contributions

Conceptualization: AARM, GTB, JAC; Data Curation: AFB, FRRM, JAC; Formal analysis: FRRM, GLW, JAC, PEB; Funding acquisition: AFB, BAC, VSS; Investigation: AARM, ARR, DCO, GTB, JAC; Methodology: AFB, FR, GLW, MANV, PEB, RFCS, RGS, VSS; Project administration: AFB, BAC, JAC; Resources: RFCS, RGS; Software: GLW; Supervision: AFB, JAC; Visualization: AFB, FRRM, GLW, JAC; Writing (Original Draft): AARM, GTB, JAC; Writing (Review & Editing): AARM, AFB, ARR, BAC, DCO, FR, FRRM, GLW, GTB, JAC, MANV, PEB, RFCS, RGS, VSS.

## Data availability

The raw data were submitted to the National Center of Biotechnology Information (NCBI) under BioProject PRJNA1205703, BioSamples from SAMN46054717 to SAMN46054740; from SAMN46350146 to SAMN46350421; from SRA SRR31872231 to SRR31872254; and from SRR32069178 to SRR32069453. The sequences have been submitted to Genbank/NCBI, and can be found with the accession numbers PQ650654.2, PQ963794.2, PV975644-PV975703.

## References

Almeida, Marilene F., Adriana R. Rosa, Miriam M. Sodré, Luzia F. A. Martorelli, and José T. Netto. 2015. “Fauna de Morcegos (Mammalia, Chiroptera) E a Ocorrência de Vírus Da Raiva Na Cidade de São Paulo, Brasil.” Veterinaria E Zootecnia 22 (1): 89–100.

Anderson, Marti J. 2008. “A New Method for Non-Parametric Multivariate Analysis of Variance.” Austral Ecology 26 (1): 32–46.

Asplund, M., K. R. Kjartansdóttir, S. Mollerup, et al. 2019. “Contaminating Viral Sequences in High-Throughput Sequencing Viromics: A Linkage Study of 700 Sequencing Libraries.” Clinical Microbiology and Infection: The Official Publication of the European Society of Clinical Microbiology and Infectious Diseases 25 (10): 1277–1285.

Bazzoni, Emanuela, Carla Cacciotto, Rosanna Zobba, Marco Pittau, Vito Martella, and Alberto Alberti. 2024. “Bat Ecology and Microbiome of the Gut: A Narrative Review of Associated Potentials in Emerging and Zoonotic Diseases.” Animals: An Open Access Journal from MDPI 14 (20). 10.3390/ani14203043.

Bentim Góes, Luiz Gustavo, Carlo Fischer, Angélica Cristine Almeida Campos, et al. 2022. “Highly Diverse Arenaviruses in Neotropical Bats, Brazil.” Emerging Infectious Diseases 28 (12): 2528–2533.

Branda, Francesco, Grazia Pavia, Alessandra Ciccozzi, et al. 2024. “Zoonotic Paramyxoviruses: Evolution, Ecology, and Public Health Strategies in a Changing World.” Viruses 16 (11). 10.3390/v16111688.

Buchfink, Benjamin, Klaus Reuter, and Hajk-Georg Drost. 2021. “Sensitive Protein Alignments at Tree-of-Life Scale Using DIAMOND.” Nature Methods 18 (4): 366–368.

Capella-Gutiérrez, Salvador, José M. Silla-Martínez, and Toni Gabaldón. 2009. “trimAl: A Tool for Automated Alignment Trimming in Large-Scale Phylogenetic Analyses.” Bioinformatics (Oxford, England) 25 (15): 1972–1973.

Cerri, Agustina, Elisa M. Bolatti, Tomaz M. Zorec, et al. 2023. “Identification and Characterization of Novel Alphacoronaviruses in Tadarida Brasiliensis (Chiroptera, Molossidae) from Argentina: Insights into Recombination as a Mechanism Favoring Bat Coronavirus Cross-Species Transmission.” Microbiology Spectrum, September 11, e0204723.

Claro, Ingra M., Mariana S. Ramundo, Thais M. Coletti, et al. 2021. “Rapid Viral Metagenomics Using SMART-9N Amplification and Nanopore Sequencing.” Wellcome Open Research 6 (September): 241.

Cláudio Vinícius C., Roberto L. M. Novaes, Alfred L. Gardner, et al. 2023. “Taxonomic Re-Evaluation of New World Eptesicus and Histiotus (Chiroptera: Vespertilionidae), with the Description of a New Genus.” Zoologia (Curitiba, Brazil) 40 (e22029). 10.1590/s1984-4689.v40.e22029.

Diaz, M. M., R. Solari, R. Gregorin, L. F. Aguirre, and R. M. Barquez. 2021. Clave de Identificación de Los Murciélagos Neotropicales. Vol. 207. Tucumán, Argentina.

Edgar, Robert C. 2004. “MUSCLE: Multiple Sequence Alignment with High Accuracy and High Throughput.” Nucleic Acids Research 32 (5): 1792–1797.

Festa, Francesca, Leonardo Ancillotto, Luca Santini, et al. 2023. “Bat Responses to Climate Change: A Systematic Review.” Biological Reviews of the Cambridge Philosophical Society 98 (1): 19–33.

Garbino, Guilherme S. T., Vinícius C. Cláudio, Renato Gregorin, et al. 2024. “Updated Checklist of Bats (Mammalia: Chiroptera) from Brazil.” Zoologia (Curitiba, Brazil) 41. 10.1590/s1984-4689.v41.e23073.

Geoghegan, Jemma L., Francesca Di Giallonardo, Michelle Wille, et al. 2021. “Virome Composition in Marine Fish Revealed by Meta-Transcriptomics.” Virus Evolution 7 (1): veab005.

Gilbert, Ruth, and Susan J. Cliffe. 2016. “Public Health Surveillance.” In Public Health Intelligence. Springer International Publishing.

Guindon, Stéphane, Jean-François Dufayard, Vincent Lefort, Maria Anisimova, Wim Hordijk, and Olivier Gascuel. 2010. “New Algorithms and Methods to Estimate Maximum-Likelihood Phylogenies: Assessing the Performance of PhyML 3.0.” Systematic Biology 59 (3): 307–321.

He, Xiaozhou, Xu Wang, Guohao Fan, et al. 2022. “Metagenomic Analysis of Viromes in Tissues of Wild Qinghai Vole from the Eastern Tibetan Plateau.” Scientific Reports 12 (1). 10.1038/s41598-022-22134-y.

Ikegame, Satoshi, Jillian C. Carmichael, Heather Wells, et al. 2023. “Metagenomics-Enabled Reverse-Genetics Assembly and Characterization of Myotis Bat Morbillivirus.” Nature Microbiology 8 (6): 1108–1122.

Kalyaanamoorthy, Subha, Bui Quang Minh, Thomas K. F. Wong, Arndt von Haeseler, and Lars S. Jermiin. 2017. “ModelFinder: Fast Model Selection for Accurate Phylogenetic Estimates.” Nature Methods 14 (6): 587–589.

Kawasaki, Junna, Keizo Tomonaga, and Masayuki Horie. 2023. “Large-Scale Investigation of Zoonotic Viruses in the Era of High-Throughput Sequencing.” Microbiology and Immunology 67 (1): 1–13.

Letko, Michael, Stephanie N. Seifert, Kevin J. Olival, Raina K. Plowright, and Vincent J. Munster. 2020. “Bat-Borne Virus Diversity, Spillover and Emergence.” Nature Reviews. Microbiology 18 (8): 461–471.

Li, Heng. 2018. “Minimap2: Pairwise Alignment for Nucleotide Sequences.” Bioinformatics (Oxford, England) 34 (18): 3094–3100.

Li, Heng, Bob Handsaker, Alec Wysoker, et al. 2009. “The Sequence Alignment/Map Format and SAMtools.” Bioinformatics (Oxford, England) 25 (16): 2078–2079.

Luz, Hermes Ribeiro, Sebastián Muñoz-Leal, Juliana Cardoso de Almeida, João Luiz Horacio Faccini, and Marcelo Bahia Labruna. 2016. “Ticks Parasitizing Bats (Mammalia: Chiroptera) in the Caatinga Biome, Brazil.” Revista Brasileira de Parasitologia Veterinaria [Brazilian Journal of Veterinary Parasitology] 25 (4): 484–491.

Mackenzie, John S., James E. Childs, Hume E. Field, Lin-Fa Wang, and Andrew C. Breed. 2016. “The Role of Bats as Reservoir Hosts of Emerging Neuroviruses.” In Neurotropic Viral Infections. Springer International Publishing.

Miller, Ruth R., Vincent Montoya, Jennifer L. Gardy, David M. Patrick, and Patrick Tang. 2013. “Metagenomics for Pathogen Detection in Public Health.” Genome Medicine 5 (9): 81.

Minh, Bui Quang, Heiko A. Schmidt, Olga Chernomor, et al. 2020. “IQ-TREE 2: New Models and Efficient Methods for Phylogenetic Inference in the Genomic Era.” Molecular Biology and Evolution 37 (5): 1530–1534.

Minter, Amanda, Graham F. Medley, and T. Déirdre Hollingsworth. 2024. “Using Passive Surveillance to Maintain Elimination as a Public Health Problem for Neglected Tropical Diseases: A Model-Based Exploration.” Clinical Infectious Diseases: An Official Publication of the Infectious Diseases Society of America 78 (Supplement_2): S169–S174.

Nunes, Hannah, Fabiana Lopes Rocha, and Pedro Cordeiro-Estrela. 2017. “Bats in Urban Areas of Brazil: Roosts, Food Resources and Parasites in Disturbed Environments.” Urban Ecosystems 20 (4): 953–969.

PAHO. 2025. “Instituto Todos pela Saúde, OPAS/OMS, Ministério da Saúde do Brasil e parceiros promovem oficina sobre vigilância em morcegos como estratégia de detecção de patógenos.” February 11. https://www.paho.org/pt/noticias/11-2-2025-instituto-todos-pela-saude-opasoms-ministerio-da-saude-do-brasil-e-parceiros.

PAHO. 2026. “Relatório da oficina técnica sobre o uso da vigilância em morcegos como estratégia de detecção de patógenos com potencial epidêmico e pandêmico.” March 9. https://www.paho.org/pt/documentos/relatorio-da-oficina-tecnica-sobre-uso-da-vigilancia-em-morcegos-como-estrategia.

Quan, Phenix-Lan, Cadhla Firth, Juliette M. Conte, et al. 2013. “Bats Are a Major Natural Reservoir for Hepaciviruses and Pegiviruses.” Proceedings of the National Academy of Sciences of the United States of America 110 (20): 8194–8199.

Quinlan, Aaron R., and Ira M. Hall. 2010. “BEDTools: A Flexible Suite of Utilities for Comparing Genomic Features.” Bioinformatics (Oxford, England) 26 (6): 841–842.

Rahimian, Mohammadreza, and Bahman Panahi. 2024. “Metagenome Sequence Data Mining for Viral Interaction Studies: Review on Progress and Prospects.” Virus Research 349 (199450): 199450.

Reis, Nelio R. dos, Adriano L. Peracchi, Isaac P. de Lima, and Wagner A. Pedro. 2006. “Riqueza de Espécies de Morcegos (Mammalia, Chiroptera) Em Dois Diferentes Habitats, Na Região Centro-Sul Do Paraná, Sul Do Brasil.” Revista Brasileira de Zoologia 23 (3): 813–816.

Sasaya, Takahide, Gustavo Palacios, Thomas Briese, et al. 2023. “ICTV Virus Taxonomy Profile: Phenuiviridae 2023.” The Journal of General Virology 104 (9). 10.1099/jgv.0.001893.

Sayler, Katherine A., Anthony F. Barbet, Casey Chamberlain, et al. 2014. “Isolation of Tacaribe Virus, a Caribbean Arenavirus, from Host-Seeking Amblyomma Americanum Ticks in Florida.” PloS One 9 (12): e115769.

Schaeffer, Reagan, Gun Temeeyasen, and Ben M. Hause. 2022. “Alphacoronaviruses Are Common in Bats in the Upper Midwestern United States.” Viruses 14 (2): 184.

Temmam, Sarah, Bernard Davoust, Jean-Michel Berenger, Didier Raoult, and Christelle Desnues. 2014. “Viral Metagenomics on Animals as a Tool for the Detection of Zoonoses prior to Human Infection?” International Journal of Molecular Sciences 15 (6): 10377–10397.

Van Brussel, Kate, and Edward C. Holmes. 2022. “Zoonotic Disease and Virome Diversity in Bats.” Current Opinion in Virology 52 (February): 192–202.

Vizotto, L. D., and V. A. Taddei. 1973. Chave Para Determinação de Quirópteros Brasileiros. São José do Rio Preto, Gráfica Francal.

Wallau, Gabriel da Luz, Eder Barbier, Lais Ceschini Machado, et al. 2025. “Ambecovirus, a Novel Betacoronavirus Subgenus Circulating in Neotropical Bats, Sheds New Light on Bat-Borne Coronaviruses Evolution.” Virus Evolution 11 (1): veaf094.

Wallau, Gabriel Luz, Eder Barbier, Alexandru Tomazatos, Jonas Schmidt-Chanasit, and Enrico Bernard. 2023. “The Virome of Bats Inhabiting Brazilian Biomes: Knowledge Gaps and Biases towards Zoonotic Viruses.” Microbiology Spectrum 11 (1): e0407722.

Wells, Heather L., Elizabeth Loh, Alessandra Nava, et al. 2022. “Classification of New Morbillivirus and Jeilongvirus Sequences from Bats Sampled in Brazil and Malaysia.” Archives of Virology 167 (10): 1977–1987.

WHO. 2024. “Pathogens Prioritization: A Scientific Framework for Epidemic and Pandemic Research Preparedness.” https://www.who.int/publications/m/item/pathogens-prioritization-a-scientific-framework-for-epidemic-and-pandemic-research-preparedness.

Wu, Zhiqiang, Yelin Han, Bo Liu, et al. 2021. “Decoding the RNA Viromes in Rodent Lungs Provides New Insight into the Origin and Evolutionary Patterns of Rodent-Borne Pathogens in Mainland Southeast Asia.” Microbiome 9 (1): 18.

Xu, Ziqian, Yun Feng, Xinxin Chen, et al. 2022. “Virome of Bat-Infesting Arthropods: Highly Divergent Viruses in Different Vectors.” Journal of Virology 96 (4): e0146421.

