## Supplementary Methods 1 for "Viral metagenomics of synanthropic urban bats: a surveillance strategy for uncovering potentially zoonotic viruses"

### Selection of bat specimens for metagenomic sequencing

#### Objective

To prioritize bat specimens collected via passive rabies surveillance for nucleic acid extraction and metagenomic sequencing, with the goal of maximizing taxonomic, temporal, and geographic diversity, and the likelihood of detecting zoonotic viruses.

#### Materials and Data Requirements

- Intake form with complete metadata: collection date, location, species identification, age class and circumstances of collection.
- Reference data: historical routine collection data and sequencing frequencies (per species and municipalities).

#### Procedure

##### Step 1: Initial screening – apply exclusion criteria

Assess tissue integrity:

- If the target tissue is intact (no signs of autolysis or decomposition), proceed.
- If degraded, **exclude** specimen.

Determine age group:

- Include only adult or juvenile specimens.
- If identified as infant/neonate, **exclude** specimen.

Check rabies diagnostics:

- Include rabies negative specimens.
- If positive, **exclude** specimen.

***Note****: Excluded specimens may still be used for other studies, but are not eligible for metagenomics.*

##### Step 2: Metadata verification

Confirm availability of the following minimal metadata:

- Date of collection
- Collection location (e.g., municipality)
- Taxonomic identification (genus and species)

##### Step 3: Assign inclusion scores (0 or 1 point per criterion)

Evaluate the following binary variables. Assign 1 point for "Yes", 0 points for "No".

| **Criteria** | **Criterion** | **Scoring question** |
| --- | --- | --- |
| 1 | Metadata completeness | Is minimal metadata available? |
| 2 | Atypical event | Was the bat collected under atypical conditions (e.g., indoors, fallen, active during the day)? |
| 3 | Human/animal contact | Was there confirmed or likely contact with humans or domestic animals? |
| 4 | Undersampled location | Is the specimen from a location contributing <15% of total sequenced bats? |
| 5 | Rare species | Does the species represent <1% of all collected bats (based on routine collection data)? |
| 6 | Undersampled species | Has the species contributed <15% of all sequenced bats so far? |
| 7 | Undersampled time point | Is the collection month below the 10% threshold for temporal sampling balance? |

##### Step 4: Prioritization Based on Final Score

Calculate total inclusion score (sum of IN1–IN7):

- If **score ≥ 4**, specimen is **eligible** for metagenomic sequencing.
- If **score < 4**, specimen is **excluded** from sequencing.

***Note****: Specimens scoring 5–7 are prioritized for sequencing to ensure diversity and high potential for viral detection.*

#### Notes

1. Scoring thresholds for species and municipality representation were dynamically updated as new metadata became available.
2. A decision tree summarizing this protocol is shown in the main manuscript in Figure 1.
