## Supplementary Methods 2 for "Viral metagenomics of synanthropic urban bats: a surveillance strategy for uncovering potentially zoonotic viruses"

Sensitivity analysis with three *de novo* assemblers

Before conducting the full metagenomic analysis workflow, we performed a preliminary sensitivity assessment using three different de novo assemblers: Flye 2.9.6 (Kolmogorov et al., 2019), Raven v1.8.3 (Vaser & Šikić, 2021) , and MEGAHIT v1.2.9 (Li et al., 2015). Flye was prioritized for initial testing due to its optimization for nanopore sequencing data, which is characterized by long and noisy reads. However, the results were limited: only 442 viral contigs were recovered, spanning nine viral families—none of which were of zoonotic relevance. Raven yielded even fewer results, with just 30 contigs associated with six viral families, all of which were bacteriophages. In contrast, MEGAHIT produced a markedly richer assembly, recovering 4,285 viral contigs from 43 viral families. Among these, nine families were of public health interest, and all viruses subjected to further detailed analyses in this study were identified through the MEGAHIT assembly.

A review of the documentation for Flye and Raven suggests that their underperformance stems from algorithmic assumptions tailored for long reads. For example, Flye requires a minimum overlap of 1,000 nucleotides (nt) between reads for contig assembly. Given that our nanopore libraries were fragmented—with a median contig size of approximately 350 nt—these parameters rendered both Flye and Raven ineffective for this dataset. MEGAHIT, in contrast, does not impose such constraints and was originally designed for assembling short reads, making it more suitable for our data.

To address the potential limitations of using a short-read assembler on nanopore data, all contigs produced by MEGAHIT were subsequently polished using Medaka v2.0.3 (<https://github.com/nanoporetech/medaka>, last accessed June 23rd, 2025), a neural network-based tool developed by Oxford Nanopore Technologies that corrects sequencing errors in long-read assemblies. Additionally, viral contigs were validated via blastX (Camacho et al., 2009) searches against the NCBI non-redundant (nr) protein database. Downstream analyses—including phylogenetic reconstruction and protein structural modeling—focused on the translated products of predicted viral open reading frames.

In summary, our sensitivity analysis showed that a pipeline combining MEGAHIT (a short-read assembler) with post-assembly polishing by Medaka was the most effective strategy for identifying viral contigs and characterizing viruses with zoonotic potential in this study. We anticipate that this finding will generalize to other datasets generated using the SMART-9N protocol (Claro et al. 2023) or other nanopore-based workflows that yield relatively short reads (e.g., <1,000 nt).

**References**

1. Kolmogorov, M., Yuan, J., Lin, Y., & Pevzner, P. A. (2019). Assembly of long, error-prone reads using repeat graphs. *Nature Biotechnology*, 37(5), 540–546. https://doi.org/10.1038/s41587-019-0072-8
2. Vaser, R., & Šikić, M. (2021). Raven: a de novo genome assembler for long reads. *Bioinformatics*, 37(3), 409–415. https://doi.org/10.1093/bioinformatics/btaa654
3. Li, D., Liu, C. M., Luo, R., Sadakane, K., & Lam, T. W. (2015). MEGAHIT: an ultra-fast single-node solution for large and complex metagenomics assembly via succinct de Bruijn graph. *Bioinformatics*, 31(10), 1674–1676. https://doi.org/10.1093/bioinformatics/btv033
4. Camacho, C. et al. (2009). BLAST+: architecture and applications. BMC *Bioinformatics*, 10, 421. <https://doi.org/10.1186/1471-2105-10-421>
5. Claro, I.M. et al. (2023). Rapid viral metagenomics using SMART-9N amplification and nanopore sequencing. *Wellcome Open Research*, 6, p.241.
